# Analyzing RNA-Seq Data from *Chlamydia* with Super Broad Transcriptomic Activation: Challenges, Solutions, and Implications for Other Systems

**DOI:** 10.1101/2024.05.16.594566

**Authors:** Danny Wan, Andrew Cheng, Yuxuan Wang, Guangming Zhong, Wei Vivian Li, Huizhou Fan

## Abstract

**Motivation:** RNA sequencing (RNA-Seq) offers profound insights into the complex transcriptomes of diverse biological systems. However, standard differential expression analysis pipelines based on DESeq2 and edgeR encounter challenges when applied to the immediate early transcriptomes of *Chlamydia* spp., obligate intracellular bacteria. These challenges arise from their reliance on assumptions that do not hold in scenarios characterized by extensive transcriptomic activation and limited repression. Standard analyses using unique chlamydial RNA-Seq reads alone identify nearly 300 upregulated and about 300 downregulated genes, significantly deviating from actual RNA-Seq read trends.

**Results:** By incorporating both chlamydial and host reads or adjusting for total sequencing depth, the revised normalization methods each detected over 700 upregulated genes and 30 or fewer downregulated genes, closely aligned with observed RNA-Seq data. Further validation through qRT-PCR analysis confirmed the effectiveness of these adjusted approaches in capturing the true extent of transcriptomic activation during the immediate early phase of chlamydial infection. While the strategies employed are developed in the context of *Chlamydia*, the principles of flexible and context-aware normalization may inform adjustments in other systems with unbalanced gene expression dynamics, such as bacterial spore germination.

**Availability and implementation:** The code for reproducing the presented bioinformatic analysis is available at https://zenodo.org/records/11201379.

## Introduction

RNA sequencing (RNA-Seq) technology has revolutionized the field of molecular biology, offering unprecedented insights into the complexity of transcriptomes across a wide array of biological systems (Mortazavi, et al., 2008; Wang, et al., 2009). This high-throughput technology not only facilitates the quantification of gene expression but also enables the detection of novel transcripts, and post-transcriptional modifications. The comprehensive nature of RNA-Seq data has been instrumental in elucidating the regulatory mechanisms underpinning growth, development, and response to environmental changes.

Central to the processing and analysis of RNA-Seq data are bioinformatic pipelines. By analyzing the quality of sequencing data, aligning sequencing reads to reference genomes or transcriptomes, quantifying gene or transcript expression levels, and performing differential expression analysis, these tools enable researchers to decipher complex transcriptomic data and identify gene expression patterns critical for biological functions (Conesa, et al., 2016; Mortazavi, et al., 2008; Oshlack and Wakefield, 2009).

A critical aspect of RNA-Seq analysis pipelines is data normalization (Bullard, et al., 2010; Dillies, et al., 2012). This process aims to distinguish true biological differences in gene expression from technical artifacts (e.g., variation in sequencing depth). By removing technical variability and adjusting for differences in sequencing depth and RNA composition, normalization allows researchers to directly compare transcript abundance without the confounding effects of sample-to-sample variations in library size.

DESeq2 and edgeR are the two most-widely used bioinformatics tools designed for analyzing differential gene expression from RNA-Seq data (Love, et al., 2014; Robinson, et al., 2009). They employ size factor normalization and trimmed mean of M-values (TMM) normalization, respectively, to adjust for library size differences and RNA composition. Both methods assume that, across different conditions, the differential expression of the majority of genes is either negligible or symmetrically balanced between up- and down-regulation, which is true in most biological systems.

*Chlamydia* is an obligate intracellular bacterium that undergoes a unique developmental cycle alternating between the elementary body (EB) and reticulate body (RB) (Abdelrahman and Belland, 2005). EBs, adapted for extracellular survival and host cell infection, exhibit limited metabolic activity due to non-conducive external environments. In contrast, RBs replicate within host cells, eventually producing progeny EBs (Rockey, et al., 2024). This cycle suggests that the primary EB-to-RB differentiation and the secondary RB-to-EB differentiation involve extensive transcriptomic activation and repression, respectively. Such broad activation and repression challenge the assumptions underlying standard normalization techniques used for transcriptomic research. In this study, we demonstrate that DESeq2 and edgeR analyses, when applied solely to chlamydial RNA-Seq reads, cannot accurately capture the transcriptomic changes during this developmental cycle. However, accounting for mapped host RNA-Seq reads or the whole library size for read count normalization helps mitigate this problem. These mitigation strategies are validated by real-time reverse transcription PCR (qRT-PCR) analysis and thus highlights the necessity of integrating host RNA data to accurately interpret early-stage transcriptomic dynamics in *Chlamydia*. In addition, we propose potential strategies for analyzing systems with similar transcriptomic activation or repression challenges where host RNA-Seq reads are not available, offering broader implications for research on free-living microbes and other isolated biological systems.

## Materials and Methods

### RNA-Seq dataset

The RNA-Seq dataset analyzed in this study are available under NCBI Gene Expression Omnibus accession number GSE248988 (Wurihan, et al., 2024). The detailed RNA preparation and sequencing procedures are described in Wurihan et al (Wurihan, et al., 2024).

### DESeq2 and edgeR analyses

The reads for each sample were first analyzed using FastQC (version 0.12.1) (Andrews, 2010). Trimming of adapter sequences and removal of low-quality reads were performed using Trimmomatic (version 0.38) (Bolger, et al., 2014). Short read sequences were aligned to the CtL2 434/Bu genome including the chromosome (GCF_000068585.1_ASM6858v1) and the pL2 plasmid (AM886278) using TopHat (version 2.1.1) (Trapnell, et al., 2012) and then quantified for gene expression by HTSeq (version 0.13.5) (Anders, et al., 2015) to obtain raw read counts per gene. For the second normalization approach to be introduced below, the read sequences were also aligned to the mouse genome (GCF_000001635.27) using TopHat and qualified using HTSeq.

DESeq2 (version 1.36.0) (Love, et al., 2014) and edgeR (version 3.38.4) (Robinson, et al., 2009) were used to carry out the differential expression analysis and identify differentially expression genes between 0 and 1 hpi based two criteria: adjusted *P* value (by Benjamini-Hochberg method) ≤ 0.05 and fold change ≥ 2. When RNA-Seq data normalization was performed using unique chlamydial reads alone, the default analysis pipeline in DESeq2 and edgeR was used. When normalization was performed using unique chlamydial and host reads, we replaced the library size factors in both methods with the recalculated ones using the approach below. First, we estimated the library size as the total number of unique chlamydial and host reads in each sample. Second, we calculated the library size factor of a sample as its library size divided by the geometric mean across all samples. When normalization was performed using sequencing depth, we recalculated the library size factors as the sequencing depth of a sample divided by the geometric mean of sequencing depth across samples.

### Quantitative reverse transcription real-time PCR

qRT-PCR was performed using QuantStudio 5 real-time PCR System (Thermo Fisher Bioscientific) and Luna Universal one-step qRT-PCR kit (New England BioLabs) as previously described (Wurihan, et al., 2021). *t*-tests were conducted using Microsoft Office Excel to identify differentially expressed genes from the qRT-PCR results. *P* values were adjusted for multiple comparisons by Benjamini-Hochberg procedure to control the false discovery rate (Benjamini and Hochberg, 1995).

## Results

### Discrepancy Unnormalized and Normalized Differential Expression Analyses in Immediate Early Transcriptomic Changes of *C. trachomatis*

We analyzed RNA-Seq data from cultures of *C. trachomatis*-infected mouse fibroblast cells at 0 and 1 hour post-inoculation (hpi) (Wurihan, et al., 2024). The mapped RNA-Seq read data, as summarized in Table 1, reveal a notable 6-fold increase in the percentage of uniquely mapped *C. trachomatis* reads from the total combined reads of chlamydial and mouse origins, rising from 0.9% to 5.4% from 0 to 1 hpi (Table 1). This increase translates to a 6.3-fold rise in *C. trachomatis* transcriptome coverage, from 17.1-to 107.7-fold (Table 1). Among the 980 *C. trachomatis* genes, 792 (80.8%) exhibited a ≥2-fold increase in read counts from 0 to 1 hpi, while only 33 genes (3.4%) showed a ≥2-fold decrease (Table 2). These observations signify a broad activation of the *C. trachomatis* transcriptome during the immediate early developmental phase, despite a constant number of chlamydial cells.

**Table 1.** Summary of RNA-Seq reads obtained from cultures of *Chlamydia trachomatis* (Ct)-infected mouse cells. The 945,817 bp *C. trachomatis* transcriptome was used to calculate transcriptome coverage.

|  | 0 hpi |  |  |  | 1 hpi |  |  |  |
| --- | --- | --- | --- | --- | --- | --- | --- | --- |
|  | Replicate 1 | Replicate 2 | Replicate 3 | Average | Replicate 1 | Replicate 2 | Replicate 3 | Average |
| Reads uniquely mapped to Ct genes | 261,249 | 388,612 | 319,552 | 323,138 | 1,744,834 | 1,582,744 | 2,786,959 | 2,038,179 |
| Fold Ct transcriptome coverage (depth) | 13.8 | 20.5 | 16.9 | 17.1 | 92.2 | 83.7 | 147.3 | 107.7 |
| Reads uniquely mapped to mouse genes | 29,157,294 | 36,976,293 | 35,248,060 | 33,793,882 | 35,537,308 | 30,667,452 | 40,757,064 | 35,653,941 |
| Uniquely mapped Ct reads/total reads (%) | 0.9 | 1.0 | 0.9 | 0.9 | 4.7 | 4.9 | 6.4 | 5.4 |

**Table 2.** Number of differentially expressed genes identified with direct RNA-Seq read counting or bioinformatic programs, DESeq2 or edgeR, using uniquely mapped *C. trachomatis* (Ct) reads, unique mapped Ct and host reads, or Ct reads with sequencing depth normalization. Genes with expression changes of ≥ 2-fold expression change from 0 to 1 hpi (*P* < 0.05) are defined as DEGs.

|  |  | Unique Ct reads only | Normalization with unique Ct and host reads | Normalization with sequencing depth |
| --- | --- | --- | --- | --- |
| RNA-Seq reads | Upregulated | 792 | NA | NA |
|  | Downregulated | 33 | NA | NA |
| DESeq2 | Upregulated | 305 | 717 | 718 |
|  | Downregulated | 303 | 26 | 25 |
| edgeR | Upregulated | 349 | 727 | 746 |
|  | Downregulated | 271 | 30 | 26 |

This extensive transcriptomic activation, alongside limited repression, challenges the standard assumptions underlying DESeq2 and edgeR analyses because these tools generally assume that most genes do not exhibit significant differential expression across conditions or that the numbers of upregulated and downregulated genes are balanced (Love, et al., 2014; Robinson, et al., 2009). When analyses were confined to *C. trachomatis* reads, both DESeq2 and edgeR failed to accurately capture the transcriptomic dynamics indicated by the read counts. Specifically, DESeq2 identified only 305 genes (31.1%) with ≥2-fold upregulation and 303 genes (30.9%) with downregulation from 0 to 1 hpi (Table 2). Similarly, edgeR recognized 349 upregulated genes (35.6%) and 271 downregulated genes (27.7%) (Table 2). The numbers of upregulated genes are less than half of those suggested by direct read counts, while the reported downregulated genes are nearly 10-fold more (Table 2). Notably, DESeq2 identified 228 genes as downregulated despite their read count showing a ≥2-fold increase, while edgeR reported 111 such genes. This discrepancy highlights the limitations of using DESeq2 or edgeR to analyze *C. trachomatis* RNA-Seq reads in isolation for accurately reporting transcriptomic changes from 0 to 1 hpi.

### Improved transcriptomic predictions following inclusion of host RNA-Seq reads

Given that host mouse reads constitute 95 to 99% of all uniquely mapped reads and considering that extensive changes in the host transcriptome from 0 to 1 hour post-inoculation (hpi) were not anticipated (Hayward, et al., 2021; Humphrys, et al., 2013), we hypothesized that incorporating host read counts into the normalization step of DESeq2 and edgeR analyses would yield a more accurate report of chlamydial transcriptomic changes during this period. This hypothesis was based on the premise that, despite the transcriptomic activation occurring within chlamydial cells, incorporating the vast majority of host reads enables a more precise estimation of the true library size.

By including mapped host RNA-Seq read counts in the analysis (see Methods for details), the number of genes identified as upregulated by DESeq2 rose from 305 to 717, and the count of downregulated genes sharply declined from 303 to 26. EdgeR analysis showed a similar trend, with upregulated genes increasing from 349 to 727, and downregulated genes decreasing from 271 to 30 (Table 2). These significant revisions in gene regulation patterns align closely with the read count trends observed, affirming the broad activation of the chlamydial transcriptome during the immediate early phase of infection.

### Improved transcriptomic predictions through sequencing depth normalization

As an alternative to including aligned host reads in the DESeq2 and edgeR analyses, we utilized sequencing depth (total read counts) to calculate the library size factor and perform normalization (Methods). This method does not specifically rely on uniquely mapped mouse reads and offers a computationally more efficient approach to account for the sample’s overall complexity. The DESeq2 analysis with sequencing depth normalization resulted in the identification of one additional upregulated gene and one fewer downregulated gene compared to the analysis using all uniquely mapped counts (Table 2). The edgeR analysis with sequencing depth normalization also demonstrated a high consistency with the previous analysis, revealing 19 additional upregulated genes and four fewer downregulated genes. These outcomes suggest that both strategies, using uniquely mapped *C. trachomatis* and host reads or the overall sequencing depth for read count normalization, are effective in accurately identifying differentially regulated genes.

### Validation of Broad Transcriptomic Activation by qRT-PCR Analysis

To validate the RNA-Seq findings derived from the two normalization approaches, by uniquely mapped reads or the sequencing depth, we conducted qRT-PCR analysis on a subset of *C. trachomatis* genes. Specifically, we analyzed 15 genes out of approximately 350 that appeared unchanged when analyzed using only chlamydial reads but were identified as upregulated through the two proposed normalization approaches. Additionally, we tested 5 genes that were initially categorized as downregulated in the analysis using only chlamydial reads but were recognized as upregulated upon the inclusion of mapped host reads or overall sequencing depth.

The qRT-PCR analysis revealed that 14 of the 15 genes tested exhibited more than a 2-fold upregulation from 0 to 1 hpi (with statistical significance), while the other gene displayed nearly a 1.6-fold increase (Table 3; Supplementary Table 1). Notably, none of the genes showed downregulation. These findings confirm the effectiveness of including mapped host reads and applying library size normalization in accurately identifying transcriptomic changes during the immediate early phase of *C. trachomatis* infection.

**Table 3.** Comparison of differential expression analysis based on RNA-Seq and qRT-PCR data. Displayed values are the fold expression changes calculated by DESeq2 or edgeR based on three normalization methods, using unique Ct reads only, unique Ct and host reads, or the overall sequencing depth, and the fold expression changes calculated based on qRT-PCR experiments. Positive values indicate increased expression and negative values indicate decreased expression from 0 to 1 hpi.

| only, unique Ct a | Unique Ct reads only |  | Unique Ct & host reads |  | Sequencing depth |  | qRT-PCR |
| --- | --- | --- | --- | --- | --- | --- | --- |
|  | DESeq2 | edgeR | DESeq2 | edgeR | DESeq2 | edgeR |  |
| rho | 1.56 | 1.90 | 22.21 | 20.31 | 23.88 | 23.38 | 8.44 |
| pmpI | 1.47 | 1.78 | 21.53 | 19.48 | 23.17 | 22.45 | 4.18 |
| atpI | 1.34 | 1.63 | 19.12 | 17.24 | 20.60 | 19.90 | 27.02 |
| pdhC | 1.25 | 1.51 | 18.41 | 16.14 | 19.89 | 18.84 | 7.01 |
| secG | 1.17 | 1.43 | 18.00 | 16.54 | 19.28 | 18.95 | 10.18 |
| incD | 1.01 | 1.22 | 15.42 | 13.91 | 16.58 | 16.04 | 4.08 |
| artJ | -1.08 | 1.12 | 13.33 | 12.01 | 14.36 | 13.88 | 6.95 |
| mip | -1.13 | 1.08 | 12.47 | 11.35 | 13.42 | 13.10 | 11.33 |
| pgi | -1.14 | 1.07 | 12.41 | 11.18 | 13.38 | 12.97 | 11.80 |
| aroL | -1.27 | -1.03 | 11.06 | 9.97 | 11.91 | 11.51 | 13.57 |
| ldh | -1.39 | -1.15 | 9.13 | 8.15 | 9.80 | 9.31 | 4.43 |
| ctl0271 | -1.49 | -1.21 | 9.40 | 8.58 | 10.11 | 9.88 | 4.45 |
| ihfA | -1.58 | -1.31 | 8.83 | 8.18 | 9.44 | 9.31 | 11.78 |
| murE | -1.70 | -1.42 | 7.92 | 6.80 | 8.59 | 7.99 | 6.10 |
| lpdA | -1.95 | -1.59 | 7.16 | 6.56 | 7.69 | 7.54 | 14.40 |
| pkn1 | -2.27 | -1.86 | 5.96 | 5.57 | 6.39 | 6.37 | 9.04 |
| nrdB | -2.32 | -1.90 | 6.23 | 5.57 | 6.73 | 6.48 | 33.44 |
| pknD | -3.13 | -2.58 | 4.62 | 4.20 | 4.97 | 4.84 | 15.03 |
| bioF | -4.19 | -3.45 | 3.58 | 3.19 | 3.86 | 3.71 | 2.52 |
| mdhC | -4.72 | -3.91 | 3.18 | 2.86 | 3.43 | 3.31 | 1.60 |

No change ( $|FC| < 2$ or $P > 0.05$ ) Upregulated Downregulated

## Discussion

This study underscores the critical role of context in transcriptomic analyses, particularly for obligate intracellular bacteria like *Chlamydia*. Our findings demonstrate that standard RNA-Seq normalization methods, such as those employed by DESeq2 and edgeR by default, which rely on assumptions of minimal or balanced differential expression across conditions, fall short in accurately capturing the dynamics of transcriptomic changes during the immediate early phase of *C. trachomatis* infection. The significant discrepancy between RNA-Seq counts and differential expression analyses highlights the need for methodological adjustments to account for the extensive transcriptomic activation and limited repression observed in *C. trachomatis*.

The modified normalization based on both chlamydial and host reads or the overall sequencing depth emerged as effective strategies to overcome these analytical challenges. By integrating host reads or adjusting for total sequencing depth, we were able to align the differential expression analysis more closely with the actual RNA-Seq read trends, validating the extensive activation of the chlamydial transcriptome. These approaches not only enhance the accuracy of transcriptomic predictions in the context of *C. trachomatis* infection but also offer a blueprint for analyzing other biological systems with similar challenges.

One such analogous system is the germination of bacterial spores, a process characterized by dramatic transcriptomic changes as dormant spores become metabolically active vegetative cells. Like the immediate early transcriptomic activation in *C. trachomatis*, spore germination involves a rapid and broad activation of gene expression. In transcriptomic analysis of bacterial spore germination in environmental samples, we suggest that researchers perform total sequencing depth normalization to accurately identify differentially regulated genes. Additionally, if accompanying organisms in the environment undergo significant changes, spiking-in synthetic transcripts into RNA samples prior to RNA-Seq library construction (Jiang, et al., 2011) would be helpful.

## Acknowledgements

This work was supported by grants from the National Institutes of Health (Grant # AI168279 to GZ, GM142702 to WVL, and AI154305 to HF).

